# Architecture of cell-cell junctions in situ reveals a mechanism for bacterial biofilm inhibition

**DOI:** 10.1101/2021.02.08.430230

**Authors:** Charlotte E. Melia, Jani R. Bolla, Stefan Katharios-Lanwermeyer, Daniel B. Mihaylov, Patrick C. Hoffmann, Jiandong Huo, Michael R. Wozny, Louis M. Elfari, Jan Böhning, Raymond J. Owens, Carol V. Robinson, George A. O’Toole, Tanmay A.M. Bharat

**Author notes:** **Corresponding author:** Tanmay A.M. Bharat.

## Abstract

Many bacteria, including the major human pathogen *Pseudomonas aeruginosa*, are naturally found in multicellular, antibiotic-tolerant biofilm communities, where cells are embedded in an extracellular matrix of polymeric molecules. Cell-cell interactions within *P. aeruginosa* biofilms are mediated by CdrA, a large, membrane-associated adhesin present in the extracellular matrix of biofilms, regulated by the cytoplasmic concentration of cyclic diguanylate. Here, using electron cryotomography of focused-ion beam milled specimens, we report the architecture of CdrA molecules in the extracellular matrix of *P. aeruginosa* biofilms at intact cell-cell junctions. Combining our *in situ* observations at cell-cell junctions with biochemistry, native mass spectrometry and cellular imaging, we demonstrate that CdrA forms an extended structure that projects from the outer membrane to tether cells together via polysaccharide binding partners. We go on to show the functional importance of CdrA using custom single-domain antibody (nanobody) binders. Nanobodies targeting the tip of functional cell-surface CdrA molecules could be used to inhibit bacterial biofilm formation or disrupt pre-existing biofilms in conjunction with bactericidal antibiotics. These results reveal a functional mechanism for cell-cell interactions within bacterial biofilms and highlight the promise of using inhibitors targeting biofilm cell-cell junctions to prevent or treat problematic, chronic bacterial infections.

## Introduction

Prokaryotic cells including bacteria and archaea are frequently found in nature as part of surface-attached, multicellular communities called biofilms (1-3). Biofilms constitute the majority of bacterial biomass on Earth (1, 4), representing a fundamental mode of bacterial existence. While bacterial biofilms may prove beneficial to eukaryotes as host-associated microbiomes (5, 6), the formation of pathogenic bacterial biofilms is associated with the establishment of serious, chronic, antibiotic-tolerant infections (7).

Recently, important advances have been made in understanding early events in biofilm formation (8), however the molecular mechanisms underlying how mature biofilms are formed and stablized are still poorly understood. One of the hallmarks of mature biofilms is the presence of an extracellular polymeric substance (EPS) matrix that binds bacterial cells together into a sessile community, promoting antibiotic tolerance and providing protection from other predatory organisms (9-12). The EPS matrix of biofilms is a complex mixture of molecules, consisting of proteins, polysaccharides and extracellular DNA (13). Comprehending the spatial arrangement of molecules in the EPS matrix of biofilms has been problematic (14) due to the inherent difficulty associated with high-resolution microscopic imaging inside the tissue-like environment of a biofilm. As a result, mechanisms of cellular tethering and the architecture of cell-cell junctions within biofilms are incompletely understood at the fundamental molecular level.

Nevertheless, elegant optical microscopy studies on *Vibrio cholerae* biofilms have provided clues to the internal organization of the EPS matrix, revealing that the proteins required for mature biofilm formation (RbmA, Bap1 and RbmC) fail to accumulate at the cell surface in the absence of an exopolysaccharide (called VPS) (15). *In vitro* studies of *V. cholerae* proteins have revealed an exopolysaccharide-dependent (RbmA) adhesin oligomerization pathway (16) and other studies suggest that direct interactions between the RbmA adhesin and glycans on partner cells lead to cell-cell adhesion (17). In *Escherichia coli*, an auto-aggregating adhesin known as Antigen 43 (Ag43) has been proposed to mediate cell-cell interactions in biofilms by a “Velcro-like” mechanism where two Ag43 molecules from apposing cells dimerize to bind cells to each other (18). In both of these comparatively well-characterized bacterial species, direct visualization of the EPS matrix has thus far not been performed, and the spatial arrangement of molecules at cell-cell junctions is as yet unclear.

*Pseudomonas aeruginosa* is a human pathogen of critical concern, posing a significant challenge in hospital settings due to its ability to form antibiotic-tolerant biofilms (19-21). Cell-cell interactions in the EPS matrix of *P. aeruginosa* biofilms are facilitated by the expression of a ∼220 kDa adhesin (Fig. 1A) known as CdrA, in a cyclic diguanylate (c-di-GMP) dependent manner (22). Under high cytoplasmic c-di-GMP concentrations CdrA expression is increased, and the mature CdrA protein is tethered to the outer membrane of *P. aeruginosa* cells through its membrane protein partner CdrB (22, 23). In these conditions, CdrA promotes cellular aggregation and biofilm formation by directly binding to polysaccharides in the EPS matrix of biofilms, such as the Psl or Pel polysaccharides (22, 24). When cytoplasmic c-di-GMP concentrations are lower, CdrA is cleaved and released into the extracellular milieu by the action of a periplasmic protease, promoting biofilm disaggregation (22).

**Fig 1.**
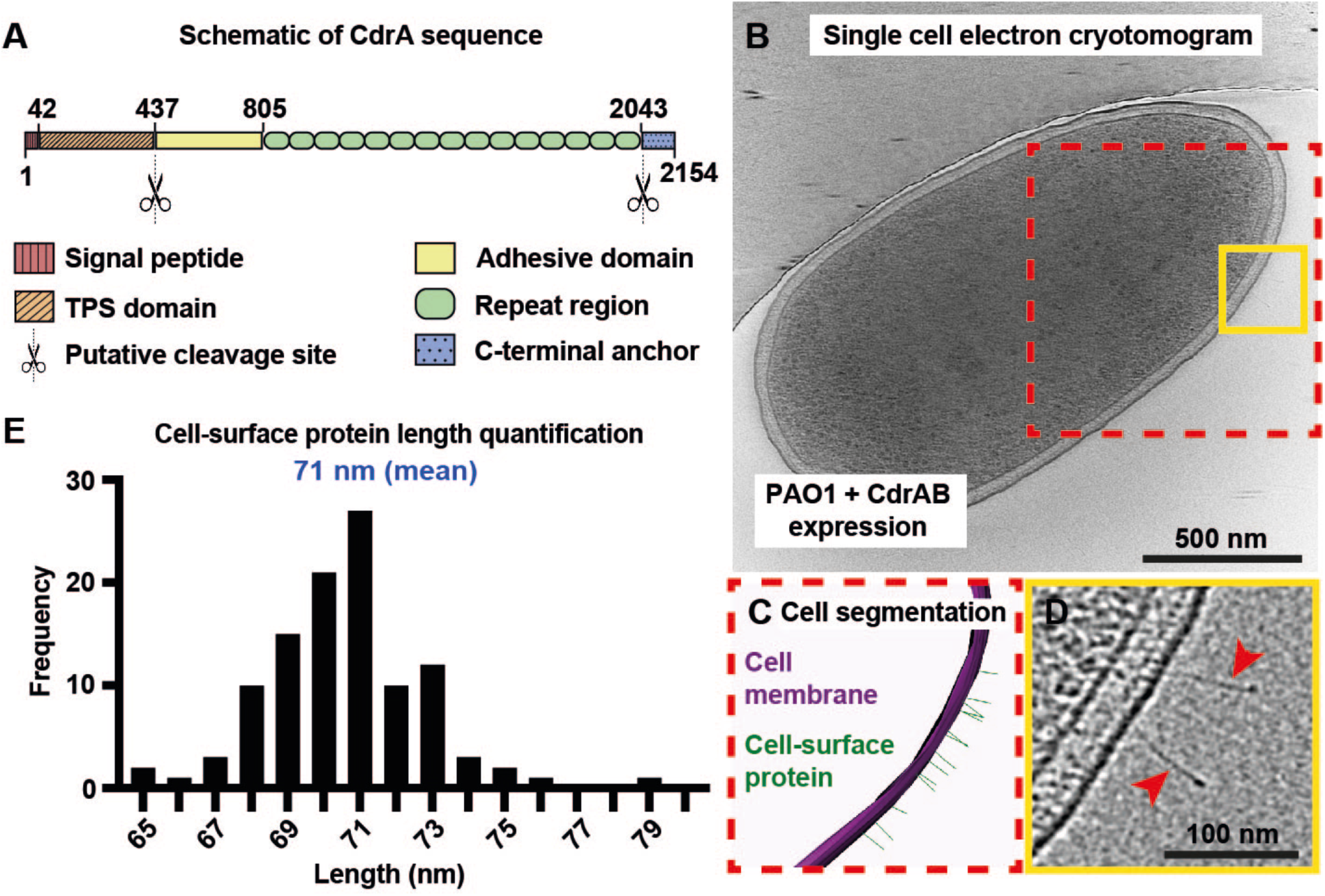
CdrAB expression results in the appearance of ∼70 nm long, matchstick-shaped protrusions on the surface of *P. aeruginosa* cells. (A) Schematic representation of the CdrA sequence highlighting previously determined and predicted functional regions including a TPS (two-partner secretion) domain, putative N-terminal cleavage site and the known C-terminal cleavage site. (B) A slice through an electron cryotomogram of a *P. aeruginosa* PAO1 cell expressing CdrAB. (C) Three-dimensional segmentation of the boxed area in (B) (red dashed line). The outer membrane of the cell (purple) and matchstick-shaped cell surface molecules (green) are shown. (D) Cropped and magnified view of the boxed region from the tomographic slice shown in (B) (solid yellow line) with matchstick-shaped protrusions indicated (red arrowheads). (E) Length quantification of cell-surface matchstick-shaped protrusions, lengths measured in electron cryotomograms (71 ± 2 nm (S.D., n=108)).

In this study, we have performed electron cryotomography (cryo-ET) of focused-ion beam milled tissue-like multicellular specimens, to image intact cell-cell junctions of *P. aeruginosa* bacteria, revealing the arrangement of CdrA in the extracellular matrix of *P. aeruginosa* biofilms. We have supplemented our *in situ* imaging with electron cryomicroscopy (cryo-EM) and native mass spectrometry (MS) experiments on biochemically purified CdrA protein, which together show that CdrA forms an extended structure at the outer membrane, forming cell-cell junctions via polysaccharide binding partners. Finally, single domain antibodies (nanobodies) raised against purified CdrA protein provided a valuable probe to test and verify all our hypotheses experimentally on wild-type bacterial biofilms, illustrating the potential for pursuing CdrA as a novel therapeutic target.

## Results

### CdrA forms an extended structure on the outer membrane of *P. aeruginosa* bacteria

To study the molecular mechanism by which CdrA tethers cells to the EPS matrix within biofilms of *P. aeruginosa*, we employed an inducible expression system, where both CdrA and CdrB are expressed. A mutation in the C-terminal part of CdrA (in the sequence TAAG, described in (23)) prevents the cleavage and release of CdrA into the extracellular environment, locking the protein in a biofilm-promoting state. Upon expression of the biofilm-promoting CdrA adhesin and its membrane anchor CdrB, *P. aeruginosa* cells form biofilm-like floccules in solution (22, 24). Cryo-ET of single cells expressing CdrAB, disassembled from cell floccules by vigorous vortexing in the presence of excess D-mannose, showed protrusions emanating from the *P. aeruginosa* cell surface (Fig. 1B-D). These ∼3 nm wide protrusions projected roughly orthogonally to the outer membrane of *P. aeruginosa* and had a broad tip, resulting in a matchstick-like appearance. The length of these matchstick-shaped protrusions was 71 nm ± 2 (S.D, n=108), as measured in three-dimensional cryo-ET data (Fig. 1E).

To establish the identity of the matchstick-shaped protrusions, molecules in the outer membrane of *P. aeruginosa* cells from the inducible CdrAB expression strain were stripped and biochemically purified (see Materials and Methods). The major component from the purification revealed a protein running at approximately 150 kDa as a single band on a gel (Fig. 2A), whose identity was confirmed to be CdrA by proteomic peptide fingerprinting. Next, electron cryomicroscopy (cryo-EM) of the MS verified, purified CdrA specimen revealed matchstick-shaped filamentous particles on the grid (Fig. 2A). A visual inspection of these particles suggested that they closely resembled the matchstick-shaped protrusions observed in whole-cell cryo-ET (Fig. 1B-D). To probe this further, cryo-ET data of the purified sample was collected and quantified, showing that the matchstick-shaped protrusions were 71 ± 1 nm (S.D., n=75) long (Fig. 2B), again indicating that they corresponded to the same cryo-EM density observed on the *P. aeruginosa* cell surface (Fig. 1B-D). Native MS of this sample showed that CdrA exists as monomers in solution (shown in Fig. 3C), indicating that the matchstick-shaped densities observed in our cryo-ET data (Fig. 1B-D) correspond to a single copy of CdrA protein projecting from the outer membrane into the extracellular environment. The measured masses of CdrA (163,286 ± 3 Da and 164,047 ± 1 Da) determined by native MS agree with previous work showing that the protein undergoes proteolytic processing into a mature functional form (22, 23).

**Fig 2.**
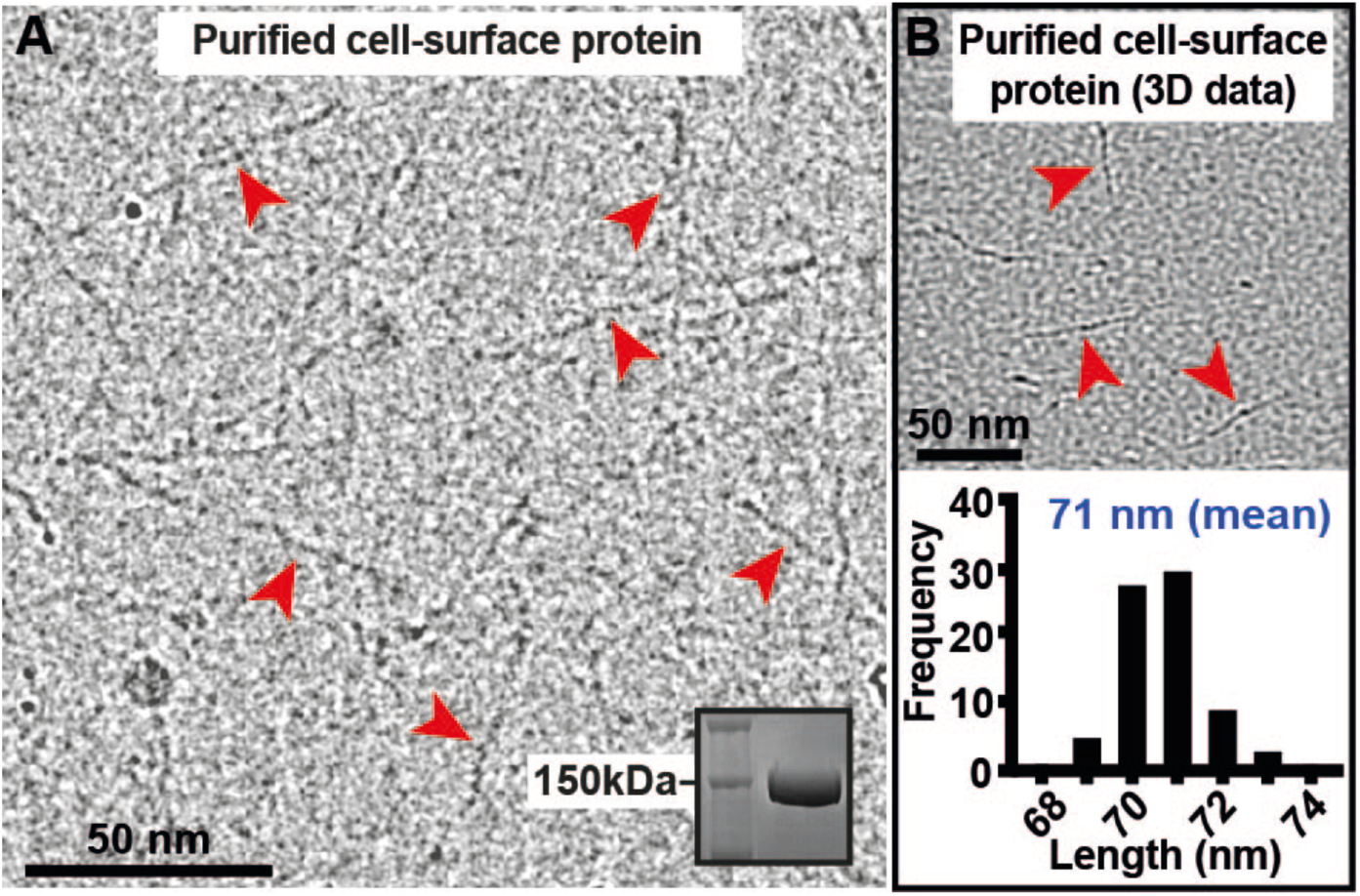
Cell-surface matchstick-shaped protrusions correspond to a mature form of CdrA. (A) Cryo-EM micrograph of protein purified from the surface of cells expressing CdrAB. Structures resembling matchstick-shaped protrusions are indicated (red arrowheads). Inset – SDS-PAGE of the purified protein. (B) Length quantification of purified protein resembling matchstick-shaped protrusions, measured in electron cryotomograms (71 ± 1 nm (S.D., n = 75)).

**Fig 3.**
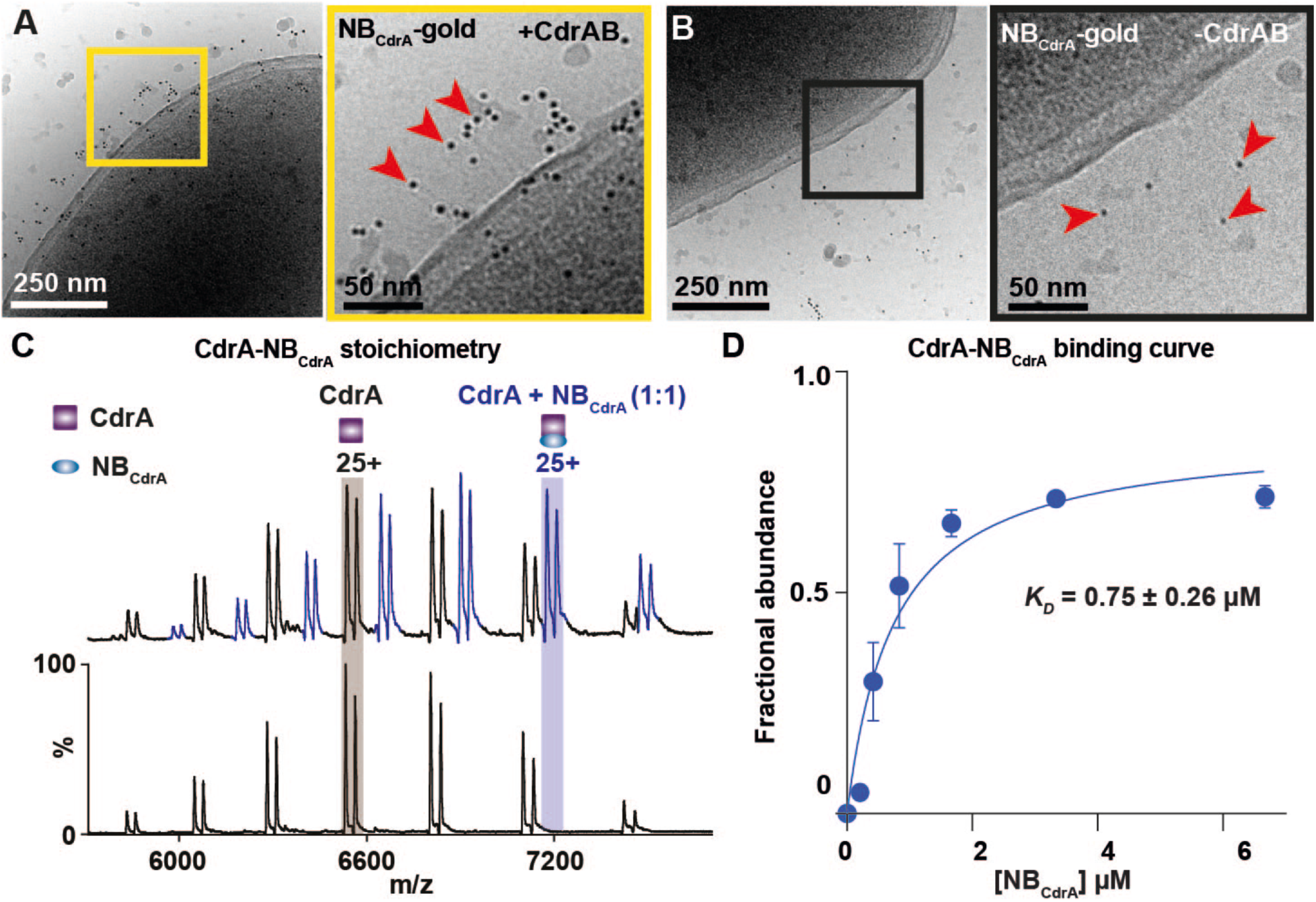
Nanobody binding to CdrA on cells and *in vitro*. (A-B), Cryo-EM micrographs of (A) cells with induced CdrAB expression or (B) control cells, labelled with a NB_CdrA_-gold conjugate (red arrowheads). Cropped and magnified views of the boxed areas in each panel are shown. (C) Native mass spectra of CdrA alone (lower spectrum) or CdrA bound to the CdrA-specific nanobody NB_CdrA_ (upper spectrum) shows binding in a 1:1 ratio. (D) A series of native MS experiments with increasing NB_CdrA_ concentrations were performed, yielding a binding curve. Each data point shows the relative fractional intensity of NB_CdrA_ binding peaks over total peak intensity (labelled as fractional abundance) versus NB_CdrA_ concentration. Standard deviation (error bars) are calculated from the average of five observed charge states in three independent experiments.

### Nanobody-targetable CdrA molecules mediate cell-cell interactions between *P. aeruginosa* bacteria

To unambiguously verify that the 71 nm matchstick-shaped protrusions observed on the *P. aeruginosa* cell surface corresponded to CdrA, we made use of single domain llama antibodies (or nanobodies), which have recently been shown to be a powerful tool for structural and cell biology (25). A panel of nanobodies was raised against purified CdrA protein, and based on a visual assessment of its ability to interfere with flocculation of *P. aeruginosa* bacteria in our inducible expression system, one positive nanobody binder was selected for further experiments. This nanobody (termed NB_CdrA_) was coupled with five-nanometer gold nanoparticles (NB_CdrA_-gold) and incubated with *P. aeruginosa* cells expressing CdrA localised to the cell surface. NB_CdrA_-gold labelling of these cells revealed a shell of gold particles specifically around cells expressing CdrAB, ∼70 nm from the cell surface (Fig. 3A). No specific NB_CdrA_-gold labelling was observed in control cells where CdrA was absent from the outer membrane (Fig. 3B), confirming that the cell-surface 71 nm matchstick-shaped protrusions correspond to CdrA molecules.

As a further verification, we performed native MS of a sample containing both purified CdrA and NB_CdrA_ and observed the formation of a 1:1 complex (Fig. 3C). To probe nanobody binding, we performed a series of native MS experiments with a constant CdrA concentration titrated against increasing NB_CdrA_ concentrations. These native experiments were used to estimate a binding affinity of NB_CdrA_ with CdrA of 0.75 ± 0.26 µM (S.D.) (Fig. 3D). These binding experiments in native MS validated our microscopic observation of NB_CdrA_-gold binding to CdrA on cells (Fig. 3A-B). Furthermore, the 1:1 binding observed in MS, together with the localization of NB_CdrA_-gold ∼70 nm from the outer membrane, strongly suggest that a region at the broad N-terminal tip of cell-surface CdrA is specifically targeted by NB_CdrA_.

We next wished to understand how CdrA mediates cell-cell interactions within the extracellular matrix of *P. aeruginosa* biofilms, by direct visualization of cell-cell junctions using high-resolution cryo-ET. We used the inducible CdrAB expression strain to promote flocculation of *P. aeruginosa* cells and deposited these floccules onto cryo-EM grids. While *P. aeruginosa* cells at the edges of the cell clump could be observed (Fig. 4A), the multicellular, tissue-like specimen was too thick for direct cryo-EM imaging. To visualize the internal arrangement of the cell-cell junctions, thin lamellae of these specimens were produced by cryo-focused ion beam (FIB) milling, which supported high-resolution imaging using cryo-ET.

**Fig 4.**
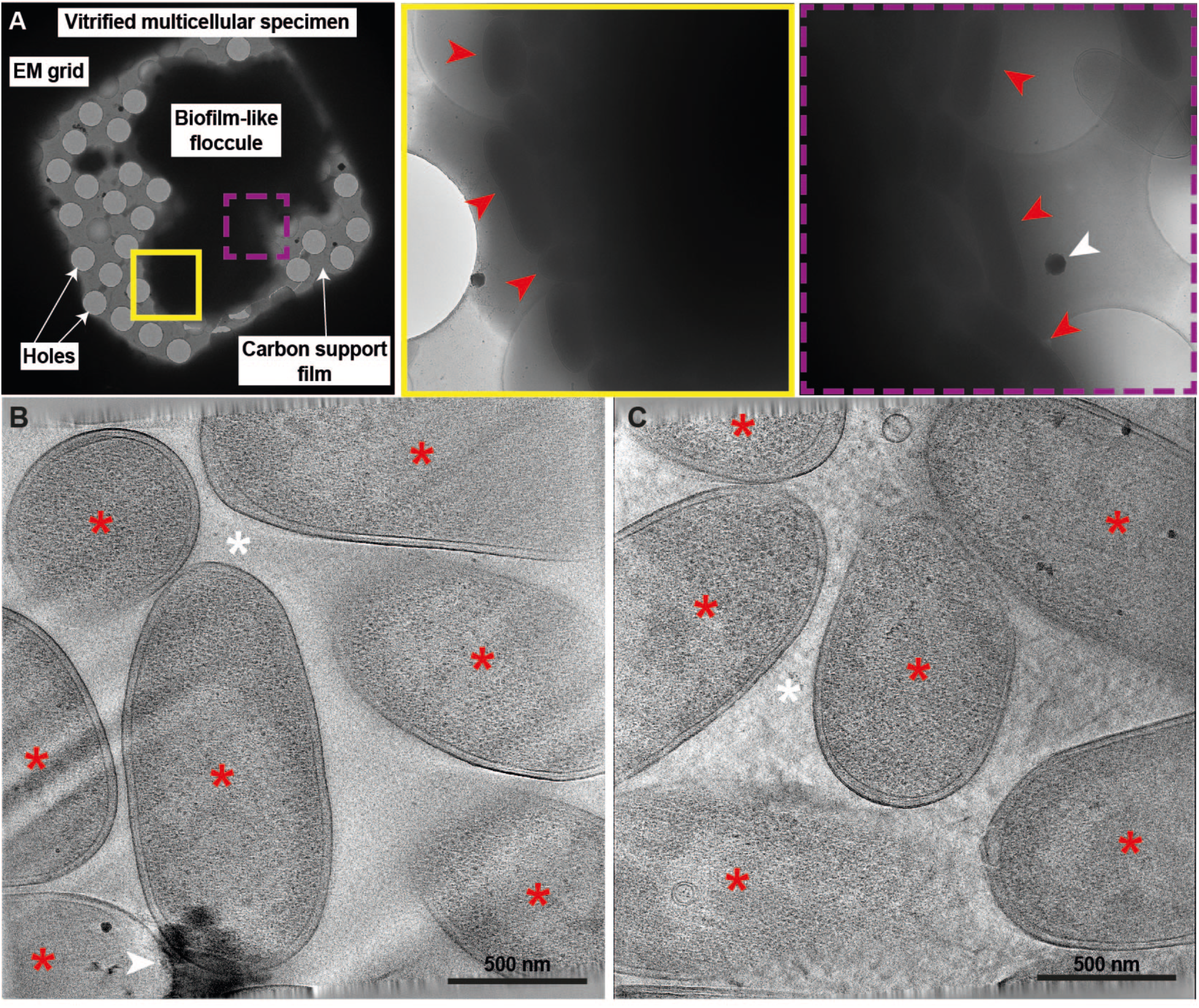
FIB-milling and cryo-ET of *P. aeruginosa* cell-cell junctions. (A) Cryo-EM of biofilm-like floccules produced by *P. aeruginosa* cells expressing CdrAB. Cryo-EM views of the multicellular aggregate. Red arrowheads indicate cells observed at the edge of the cellular aggregate (yellow solid and magenta dashed boxes). (B-C) Cryo-ET slices through FIB-milled specimens revealing cells (red asterisks) and cell-cell junctions (white asterisks). For clarity, white arrowheads indicate ice contamination.

In electron cryotomograms of cryo-FIB milled lamellae, we observed *P. aeruginosa* bacterial cells in close proximity with each other (Fig. 4B-C). A visual inspection of the cell-cell junctions between the *P. aeruginosa* cells revealed matchstick-shaped protrusions corresponding to CdrA molecules (Fig. 5A-B). These CdrA molecules were projecting outward, away from the bacterial outer membrane and extending between *P. aeruginosa* cells into the inter-cellular space (Fig. 5C). The length of these CdrA molecules was 71 ± 1 nm (S.D., n = 20), in line with the observations of CdrA on single *P. aeruginosa* cells and also of CdrA molecules after purification (Figs. 1-2, and Fig. 5D-E). In our data, direct CdrA:CdrA linkages were never observed, nor was a “Velcro-like” side-by-side configuration of CdrA molecules from apposing cells seen. The lack of direct CdrA stacking, in conjunction with the known interactions between CdrA and polysaccharides (22, 24) is consistent with a scenario in which CdrA molecules extend out of the bacterial cell surface to tether cells through interactions with polysaccharide binding partners known to be abundant in the EPS matrix.

**Fig 5.**
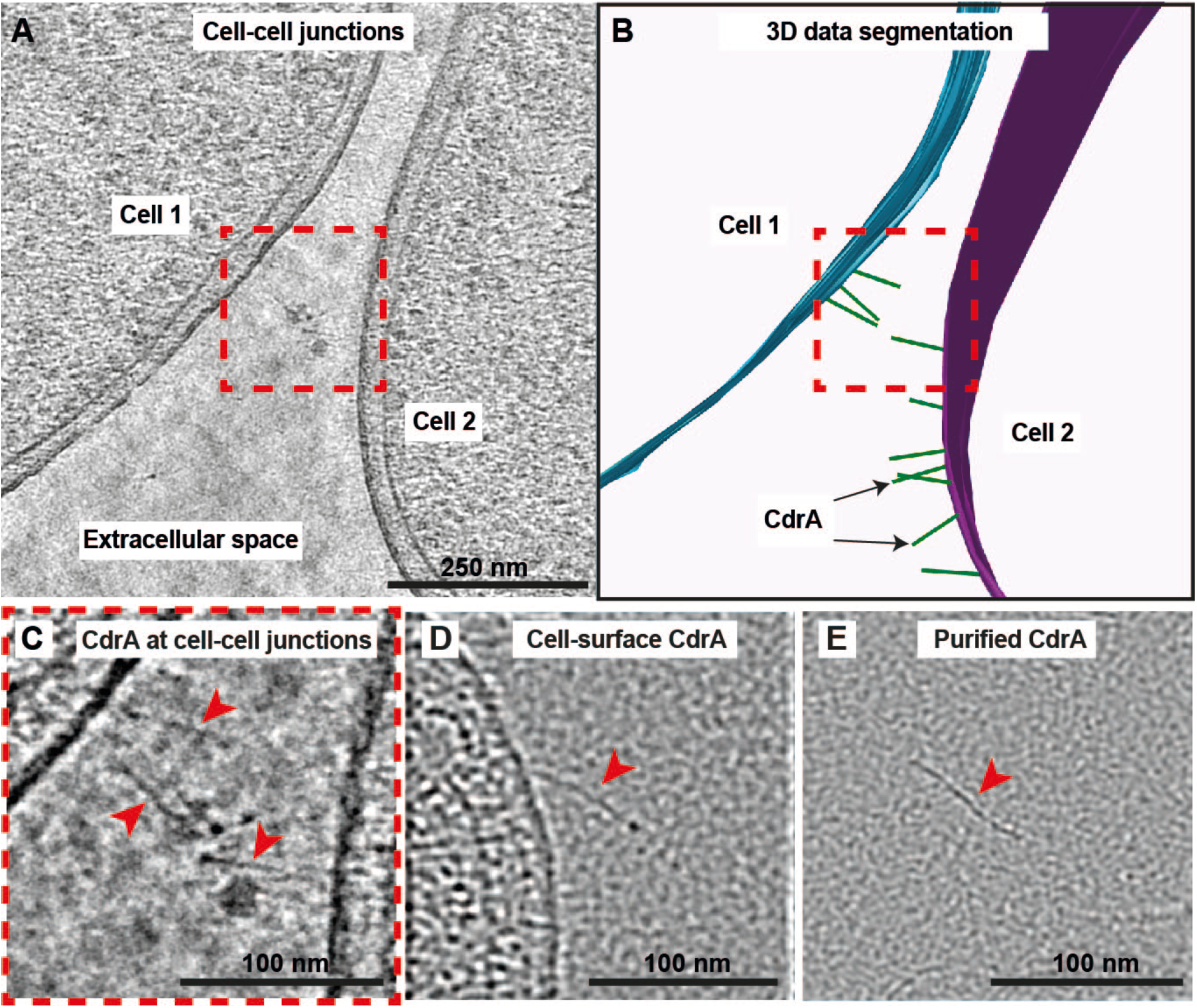
CdrA extends into the EPS matrix to mediate cell-cell interactions. (A-B), Cryo-ET slice (A) and corresponding three-dimensional segmentation (B) of a cell-cell junction within a *P. aeruginosa* PAO1 biofilm-like floccule. The multicellular specimen was processed by cryo-FIB-milling into thin lamellae suitable for high-resolution cryo-ET. Cell outer membranes (blue and purple) and CdrA (green) are highlighted in (B). (C) Enlarged view of the boxed region in (A). Comparison of CdrA at cell-cell junctions within biofilm-like floccules to (D) CdrA on the surface of single cells and (E) biochemically purified CdrA in solution. CdrA is highlighted with red arrowheads.

### The anti-CdrA nanobody (NB_CdrA_) inhibits biofilm formation and disrupts pre-formed, mature biofilms

Given the mechanistic scenario suggested by our cellular (Fig. 1), *in vitro* (Fig. 2) and *in situ* imaging (Figs. 4 and 5) where CdrA extends into and likely tethers cells to the EPS matrix, we next set out to determine whether this function of CdrA could be blocked by the targeted use of nanobodies to inhibit *P. aeruginosa* biofilm formation. We assessed whether NB_CdrA_, the nanobody shown to bind to the broad tip of CdrA molecules (Fig. 3A-B), could efficiently disrupt CdrA-mediated cell-cell adhesion and flocculation of *P. aeruginosa* using a custom microfluidics flow system. Combining the flow system with continuous fluorescence microscopy imaging, we found that the formation of *P. aeruginosa* PA14 biofilms was significantly delayed upon the addition of NB_CdrA_ (Fig. 6), showing that NB_CdrA_ can also block the function of native CdrA molecules on wild-type cells.

**Fig 6.**
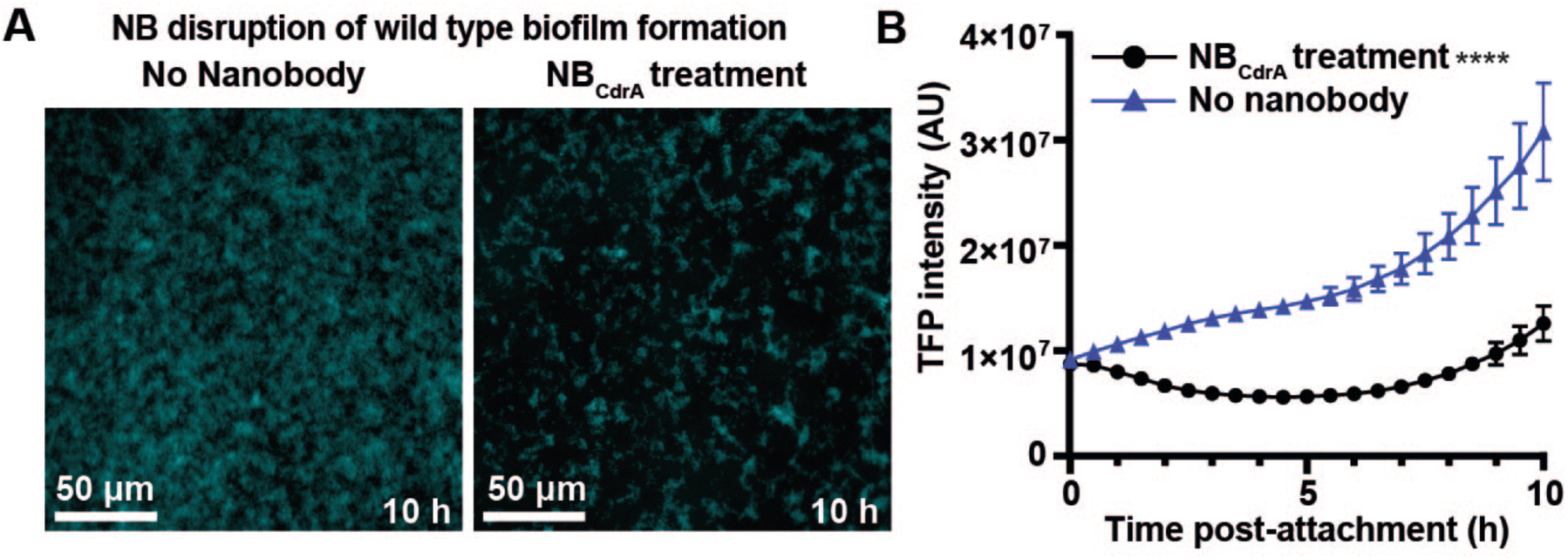
Nanobody-mediated disruption of CdrA inhibits biofilm formation. (A) Biofilms of *P. aeruginosa* PA14 expressing TFP (teal fluorescent protein) were cultivated in the presence or absence of NB_CdrA_ and monitored over 10 hours under microfluidics flow cell conditions. Representative images are shown. (B) Quantification of fluorescence in the experiments described in (A). Each time point represents three images per time point from at least three independent experiments (error bars show standard deviation). Statistical significance was assessed using the student’s t-test at all times points (****, P < 0.0001 at all times at or after 2.5 hours post treatment).

NB_CdrA_ access to cell-surface CdrA is likely to be greater in immature, developing biofilms where cell-cell junctions have not yet formed, and where diffusion deep into a multicellular specimen is not needed. We next investigated whether pre-formed, mature *P. aeruginosa* biofilms could also be disrupted by NB_CdrA_. Using the same flow setup, we found that bacterial cells in pre-formed biofilms were efficiently killed when an otherwise sub-lethal colistin antibiotic treatment was supplemented with NB_CdrA_ (Fig. 7A-B). The efficacy of bacterial killing was substantially increased when NB_CdrA_ was added earlier, during biofilm development, before the addition of colistin (Fig. 7A-B), in line with our data demonstrating the inhibitory effect of NB_CdrA_ on developing biofilms (Fig. 6). These results using wild-type *P. aeruginosa* biofilms demonstrate the key role of the CdrA protein in mediating cell-cell interactions, and highlight the importance of these interactions for effective biofilm formation, which directly promote tolerance of those biofilms to antibiotic treatment.

**Fig 7.**
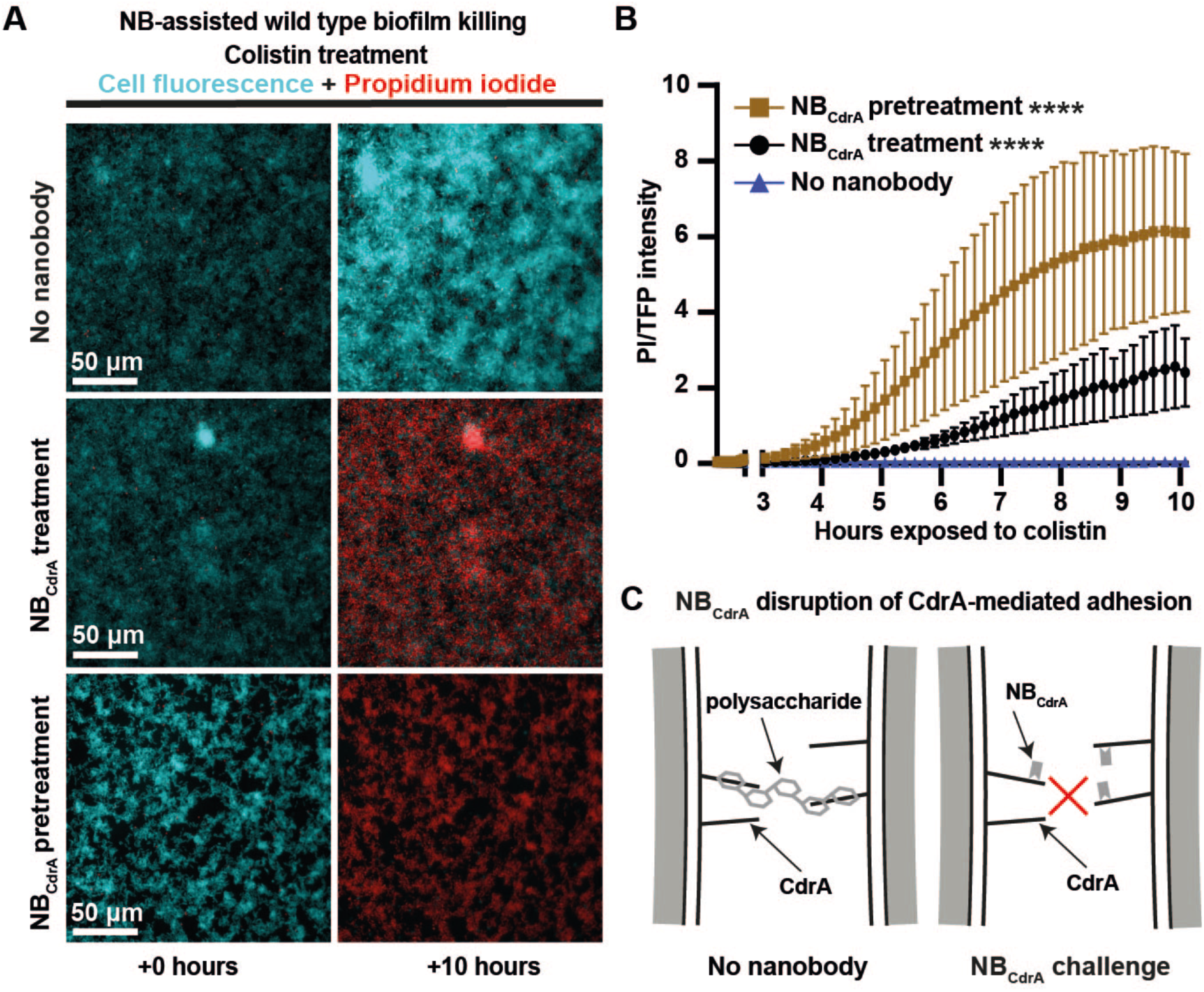
Nanobody assisted bacterial killing in pre-existing, mature biofilms. (A) Biofilms were grown in a microfluidics flow setup either in the presence (NB_CdrA_ pretreatment) or absence of NB_CdrA_ and imaged immediately after the addition of PI (propidium iodide) and a sub-lethal dose of colistin (1μg/ml) +/-NB_CdrA_ for a further ∼10 hours. Fluorescence microscopy images with TFP (blue) for live cells or propidium iodide staining (red) for dead cells are shown. (B) Quantification of fluorescence in the experiments described in (A), following the addition of colistin. The ratio of PI to TFP fluorescence was calculated over three replicate experiments (error bars denote standard deviation). Statistical significance was assessed using the student’s t-test for all time points (****, P < 0.0001, for times after 6-7 hours versus the no-nanobody control). (C) Schematic depiction of the mechanism of biofilm inhibition by targeting CdrA revealed in this study.

## Discussion

Our results establish that functional cell-surface CdrA forms an extended structure, projecting into the EPS matrix, to mediate cell-cell interactions. This arrangement is consistent with CdrA acting as a molecular tether, where copies of CdrA anchored on the cell surface at their C-termini may be glued at their N-terminal adhesive regions by secreted polysaccharides, known to be abundant in the EPS matrix of *P. aeruginosa* and previously shown to interact with CdrA (14, 24). Our results demonstrate that CdrA is a major adhesin mediating biofilm formation in wild-type PA14 *P. aeruginosa*, and shows that adhesins with similar roles, such as LecB (26), cannot effectively compensate for its disruption. While the exact contribution of different adhesins in *P. aeruginosa* biofilm development will require detailed experiments to delineate, the prominent role of CdrA-mediated cell-cell junctions is highlighted by our nanobody experiments.

This paradigm of a bacterial adhesin interacting with the EPS matrix is functionally distinct from either the alternative ‘Velcro-like’ mechanism of cell-cell adhesion proposed for *E. coli* Ag43 (18) or from the more nuanced case of *V. cholerae* where multiple adhesins and exopolysaccharides appear to regulate cell-cell interactions (15, 16). In both *E. coli* and *V. cholerae*, the spatial organisation of cell-cell junctions in biofilms has not yet been observed experimentally at high-resolution, and further research will be needed to understand the arrangement of these junctions at the molecular level. In our *in situ* cryo-ET data, while CdrA proteins were observed, extracellular polysaccharides could not be resolved. It therefore remains to be described how polysaccharides are arranged in the EPS matrix in relation to bacterial cells and other matrix molecules.

Recently, latest generation optical microscopy techniques have proven invaluable in providing novel insights into bacterial biofilm formation (4, 15, 27). In this study, we have leveraged the latest cryo-ET techniques to reveal the arrangement of cell-cell junctions that are of key importance in biofilm formation. This study highlights the utility of FIB milling and cryo-ET to deliver high-resolution insights into tissue-like multicellular specimens, which have traditionally been considered beyond the reach of structural cell biology. With increasing throughput and automation of these methods (28, 29), structure determination of molecules in cell-cell junctions may become practical, and a possible extension of this work in the future.

Moreover, we also show that biofilm formation in wild-type *P. aeruginosa* bacteria can be inhibited by targeting the filamentous CdrA adhesin with a nanobody binder that interacts with the tip of the functional protein in a 1:1 ratio. Our results are consistent with the scenario in which nanobody binding precludes the interaction between CdrA and polysaccharides in the EPS matrix (Fig. 7C). This nanobody was further shown to promote the activity of an antibiotic in killing bacterial cells in a pre-formed wild-type *P. aeruginosa* biofilm. The strategy of neutralising biofilm adhesion mechanisms may also be applicable to similar large cell-surface adhesins, such as the filamentous haemagglutinin (FHA) of *Bordetella pertussis* (30) the repeat-in-toxin (RTX) domain-containing adhesins such as SiiE of *Salmonella enterica* and LapA from *Pseudomonas fluorescens* (31-33). Such an approach may also be useful in treating bacteria whose adhesins utilise alternative proposed cell-cell adhesion mechanisms, such as Ag43 of *E. coli* (18) or RbmA of *V. cholerae* (16).

While neutralizing therapies now represent an important treatment route for many diseases, they remain relatively underexploited in the field of antimicrobials (34, 35). In the context of the increasing challenges posed by the emergence of antimicrobial resistance (36, 37), our results highlight how specific adhesins on the surface of bacterial cells may serve as promising targets for biofilm inhibition or for the prevention of chronic infections in a medical setting.

## Materials and Methods

### Cultivation of biofilm-like floccules

For CdrAB(TRRG) expression an arabinose-inducible pMQ72 plasmid system was used, as described in (23), and transformed into the PAO1 Δ*pslBCD* or PAO1 Δ*cdrA* strains, described in (22). The strains PAO1 Δ*psl* pMQ72-*cdrAB*(TRRG) and PAO1 Δ*cdrA* pMQ72-*cdrAB*(TRRG) were tested alongside a PAO1 Δ*cdra* pMQ72 empty vector control. Cultures were induced with arabinose to induce flocculation. To disaggregate cells, floccules were briefly vortexed with (for strains with wild type Psl) or without (for the Psl deletion strain) the addition of 0.5% mannose (w/v).

### CdrA protein purification

To purify CdrA, subcultures of PAO1 Δ*pslBCD* pMQ72-*cdrAB*(TRRG) were plated on LB-agar supplemented with gentamicin and arabinose to induce CdrAB expression, and incubated overnight at 37 °C. The following day, cells were scraped from the plates into PBS (phosphate buffered saline), vortexed and centrifuged to remove cells. The supernatant was polyethyleneglycol (PEG) precipitated and was centrifuged to remove contaminating proteins. The resulting soluble CdrA sample was dialysed and purified by anion exchange chromatography. Eluted fractions were analyzed by SDS-PAGE, and fractions containing CdrA were further purified by size exclusion chromatography.

### Nanobody expression and purification

To express nanobodies, nanobody phagemids were first transformed by heat shock into WK6 *E. coli* cells. Subcultures of transformants were prepared and induced with isopropyl β-D-thiogalactoside (IPTG) and incubated overnight. The following day cells were pelleted, resuspended in a lysing buffer and homogenised. His-tagged nanobodies were purified by Ni-NTA affinity chromatography and size exclusion chromatography.

### Native mass spectrometry of CdrA and nanobodies

Data were collected on a Q-Exactive UHMR mass spectrometer (ThermoFisher) and analyzed using Xcalibur 4.2 (Thermo Scientific) and UniDec (38) software packages. Nanobody binding experiments were performed by adding increasing amounts of NB_CdrA_ to a constant 2 µM CdrA. The resulting calculated binding affinity (*K*_D_) from native MS was determined by employing a nonlinear regression curve fitting for one-site specific binding in GraphPad Prism 8.0.

### Protein identification by proteomics

For protein identification, tryptic peptides were obtained by digesting the protein sample with trypsin as described in (39). Peptides separated using a chromatographic system connected to an Eclipse Tribrid Orbitrap mass spectrometer (ThermoFisher). Raw data files were processed for protein identification using MaxQuant (version 1.6.3.4) and searched against the UniProt database.

### Nanobody-gold conjugation and labelling

NB_CdrA_ was dialysed into 20 mM Tris pH 7.4, 150 mM NaCl and incubated with a ∼2.5 times molar excess of Ni-NTA 5 nm gold (Nanoprobes) for 30 minutes at room temperature. Unbound gold was separated from NB_CdrA_-gold using a PD-10 desalting column. To label cells, NB_CdrA_-gold was incubated with PAO1 Δ*cdrA* pMQ72-*cdrAB*(TRRG) floccules followed by the addition of mannose ahead of cryo-EM sample preparation.

### Microfluidics-based microscopy and analysis

To determine the effect of nanobodies on the formation of wild type (wt) biofilms, a previously described flow system was used (40). *P. aeruginosa* PA14 was visualized via fluorescence of the teal fluorescent protein (TFP). Biofilms were grown and visualized in microfluidics chambers generated through soft lithography techniques (41, 42). Bacteria were prepared as described in (43). Bacterial cultures were allowed to attach for 1 hour without flow, after which the flow rate was set to 1.0 µl/minute and imaged for 10-12 hours, as indicated.

For the biofilm-inhibition assays, the biofilm medium was supplemented either with no nanobody or 0.3 mg/ml NB_CdrA_ as indicated and imaged for ∼10 hours post inoculation. To test the impact of colistin on a preformed biofilm, bacteria were prepared, inoculated and grown in microfluidics chambers as described above. After 10 hours of allowing the biofilm to establish in the absence of any treatment, KA medium was switched with KA medium supplemented with propidium iodide (PI) and 1.0 μg/ml colistin +/-NB_CdrA,_ as indicated, and imaged for an additional ∼10 hours of incubation.

### Sample preparation for cryo-EM

Cryo-EM samples were prepared by depositing 2.5 µl of bacterial floccules, disaggregated bacterial floccules (single cells), or NB_CdrA_-gold labelled cells onto freshly glow discharged Quantifoil grids. Samples were fixed with 1% PFA applied directly on the grid, and manually blotted. Next, buffered fiducial gold was added, and the grid was blotted and plunge-frozen into liquid ethane using a Vitrobot Mark IV (ThermoFisher).

Cryo-focused ion-beam milling and scanning electron microscopy

Cryo-FIB milling of plunge-frozen biofilm-like floccules was performed as described previously (44), on a Scios DualBeam FIB/SEM microscope (FEI/ThermoFisher) equipped with a Quorum PP3010T cryo-FIB/SEM preparation system. The loading stage and milling procedure were adapted, with minor alterations, from (45). Grids were sputter-coated with platinum before milling and coated with a layer of organometallic platinum. Ion beam current for milling was reduced stepwise while adjusting the stage tilt as described in (44). Final polishing of the lamellae resulted in 150-300 nm thick lamellae.

### Transmission cryo-electron microscopy and tomography (cryo-EM and cryo-ET)

Tilt series data were collected on a Titan Krios microscope (ThermoFisher) operating at 300 kV fitted with a Quantum energy filter (Gatan) and direct electron detector (Gatan) using SerialEM software (46). A dose-symmetric tilt scheme was employed for cryo-ET data (47). Untilted video frame stacks of purified protein were collected on the same microscope using EPU software (ThemoFisher). Cryo-EM images of NB_CdrA_-gold labelled cells were collected on a Talos Arctica 200 kV cryo-TEM (ThermoFisher).

Tilt series alignment was carried out using the eTOMO graphical user interface in the IMOD software (46). CTF (contrast transfer function) parameters for the aligned stacks were estimated using CTFFIND (48) and data were reconstructed using Tomo3D (49). Contrast in cryo-ET data was enhanced via the tom_deconv deconvolution as described in (50) or via bandpass and Laplacian filtration as implemented in Fiji (51). Segmentation of image data was performed manually in IMOD (46).

## Acknowledgements

T.A.M.B. is a recipient of a Sir Henry Dale Fellowship, jointly funded by the Wellcome Trust and the Royal Society (202231/Z/16/Z). T.A.M.B would like to thank the Vallee Research Foundation, and the John Fell Fund for support. G.A.O. acknowledges support from the NIH (R37-AI83256), and C.V.R. acknowledges funding from the Medical Research Council (MR/N020413/1). J.H. is funded by the EPA Cephalosporin Fund and PPUK is supported by the Rosalind Franklin Institute EPSRC Grant no. EP/S025243/1. We thank Charlie Hitchman for assistance with the initial purification of CdrA, and Adam Costin for help with cryo-EM imaging. We also thank Carey Nadell for providing the microfluidics chambers and Wanda Kukulski for facilitating FIB milling experiments. We would like to acknowledge the MRC Laboratory of Molecular Biology Electron Microscopy Facility for access to cryo-FIB sample preparation.

